# Alterations in the intrinsic properties of striatal cholinergic interneurons after dopamine lesion and chronic L-DOPA

**DOI:** 10.1101/2020.02.14.950022

**Authors:** Se Joon Choi, Thong C. Ma, Yunmin Ding, Timothy Cheung, Neal Joshi, David Sulzer, Eugene V. Mosharov, Un Jung Kang

## Abstract

Changes in striatal cholinergic interneuron (ChI) activity are thought to contribute to Parkinson’s disease pathophysiology and dyskinesia from chronic L-3,4-dihydroxyphenylalanine (L-DOPA) treatment, but the physiological basis of these changes are unknown. We find that dopamine lesion decreases the spontaneous firing rate of ChIs, whereas chronic treatment with L-DOPA of lesioned mice increases baseline ChI firing rates to levels beyond normal activity. The effect of dopamine loss on ChIs was due to decreased currents of both hyperpolarization-activated cyclic nucleotide-gated (HCN) and small conductance calcium-activated potassium (SK) channels. L-DOPA reinstatement of dopamine normalized HCN activity, but SK current remained depressed. Pharmacological blockade of HCN and SK activities mimicked changes in firing, confirming that these channels are responsible for the molecular adaptation of ChIs to dopamine loss and chronic L-DOPA treatment. These findings suggest that targeting ChIs with channel-specific modulators may provide therapeutic approaches for alleviating L-DOPA-induced dyskinesia in PD patients.

## Introduction

The loss of dopamine (DA) neurons of the substantia nigra pars compacta results in depletion of striatal DA and the manifestation of the cardinal symptoms of Parkinson’s disease (PD). Though DA replacement therapy with its metabolic precursor L-3,4-dihydroxyphenylalanine (L-DOPA) remains the most effective treatment for PD symptoms, L-DOPA therapy can cause debilitating involuntary movements termed L-DOPA-induced dyskinesias (LID) (Fahn, 2015). Dysfunction in striatal neurocircuitry due to nigrostriatal neurodegeneration and non-physiological DA synthesis from L-DOPA by non-dopaminergic neurons are thought to underlie the development of LID. The striatum is comprised primarily of GABAergic spiny projection neurons with a small population of GABAergic and cholinergic interneurons that in mice constitute ∼5% of all striatal neurons. Though small in number, these interneurons have widespread effects on basal ganglia function (Abudukeyoumu et al., 2019; Goldberg et al., 2012) and are thought to be important contributors to both PD-related motor dysfunction and LID.

In particular, striatal cholinergic interneurons (ChIs) have been shown to contribute to movement abnormalities and LID development in rodent models of PD (Aldrin-Kirk et al., 2018; Bordia et al., 2016b; Ding et al., 2011; Divito et al., 2015; Gangarossa et al., 2016; Lim et al., 2015; Shen and Wu, 2015; Won et al., 2014). Using mice deficient for paired-like homeodomain transcription factor 3 (Pitx3) which lack substantia nigra DA neurons from birth, we previously reported that repeated L-DOPA treatment caused LID, increased baseline ChI firing rate, and potentiated ChI response to DA (Ding et al., 2011), while ablation of ChIs in a 6-hydroxydopamine (6-OHDA) model significantly attenuated LID (Won et al., 2014). Similarly, increasing ChI firing rate by chemogenetic or optogenetic methods enhanced LID expression (Aldrin-Kirk et al., 2018; Bordia et al., 2016a), while decreasing ChI firing rate by inhibiting H2 histamine receptors that are preferentially expressed in ChIs in the striatum, attenuated LID (Lim et al., 2015). Alterations in ChI activity following DA depletion have been studied, although with conflicting results (Ding et al., 2006; Maurice et al., 2015; McKinley et al., 2019; Sanchez et al., 2011; Tubert et al., 2016). However, the basis of changes in ChI physiology resulting from chronic L-DOPA treatment of DA depleted mice have not been identified.

Here, using acute striatal slice electrophysiological recordings in the dorsolateral striatum prepared from DA depleted and L-DOPA treated mice, we show that changes in intrinsic properties of ChIs cause slower spontaneous firing rates following DA depletion and faster firing rates after chronic L-DOPA treatment compared to sham controls. We found that both HCN and SK current were decreased in the DA depleted condition, whereas only SK current remained depressed after chronic L-DOPA. Changes in the activities of these two channels were sufficient to explain the observed alterations in ChI firing. Targeting altered activity of striatal ChIs with specific channel modulators may provide a potential therapeutic approach for the alleviation of LID in PD patients.

## Results

### Changes in ChI spontaneous firing rate following DA depletion and chronic L-DOPA treatment

To study alterations in ChI physiology in the parkinsonian mouse striatum, we induced dopaminergic lesions by infusing 6-OHDA unilaterally to the medial forebrain bundle (MFB). The protocol that we used produces severe and permanent lesion of both the cell bodies and axons of midbrain dopaminergic neurons (Won et al., 2014). After 3-4 weeks of recovery, the animals were randomized to receive daily IP injections of either saline (*6-OHDA* group) or L-DOPA (3 mg/kg, *chronic-LD* group) that continued for the next 3-11 weeks. Control mice (*sham* group) received vehicle MFB infusion and daily IP saline injections (**Figure 1A**). All 6-OHDA-infused mice showed severe deficits in contralateral front paw adjusting steps in the weight-supported treadmill stepping task, confirming significant lesion of the dopaminergic system. There was no difference in stepping deficit between mice assigned to the 6-OHDA or chronic-LD groups (**Figure S1A**). When challenged with a dyskinesogenic dose of L-DOPA (3 mg/kg), all lesioned mice showed abnormal involuntary movements indicative of LID, starting with the first L-DOPA injection (**Figure S1B**). Total LID magnitudes were similar between the first L-DOPA dose (representative of the 6-OHDA group) compared to mice tested after chronic L-DOPA (**Figure S1C**). However, the onset of LID was shifted to earlier time points in chronic-LD mice (**Figure S1B, D**), indicating further sensitization of LID despite maximized total LID scores from the dose of L-DOPA we used (Nadjar et al., 2009).

**Fig 1.**
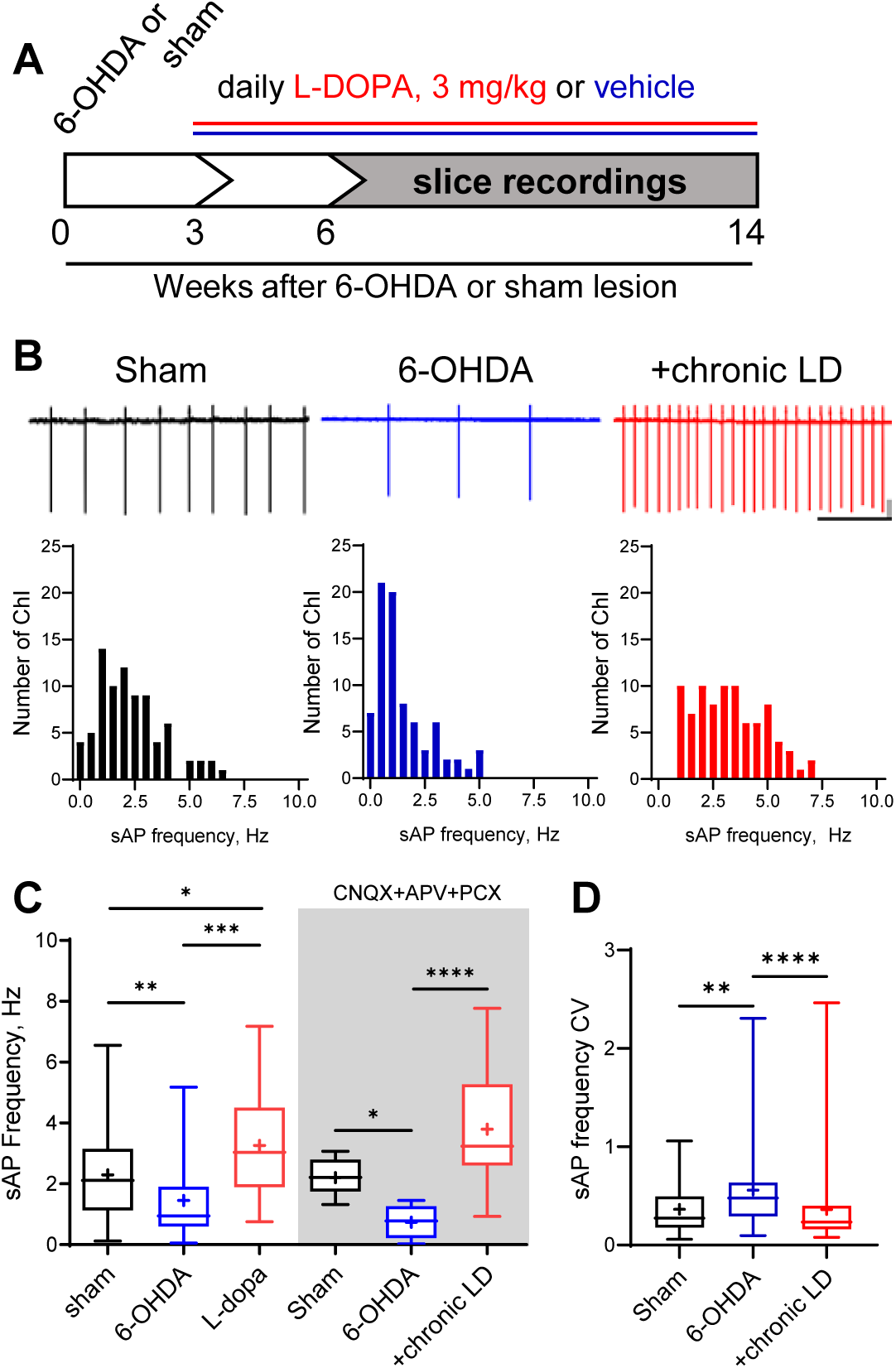
Changes in ChI spontaneous firing frequency induced by DA depletion followed by chronic L-DOPA treatment. (A) DA lesion and chronic L-DOPA treatment paradigm. 3-4 weeks after unilateral 6-OHDA lesion, mice were randomly divided into two groups to receive either saline or L-DOPA. Experimental groups included Sham: mice with vehicle injection into the MFB, 6-OHDA: mice with MFB lesions injected with daily IP saline, and chronic LD: MFB-lesioned mice treated with 3 mg/kg L-DOPA IP once daily. Electrophysiological slice recordings were carried out 3-11 weeks after the initiation of L-DOPA or saline injections. (B) Representative cell-attached recordings and distributions of average (per cell) instantaneous spontaneous action potential frequency (sAP) of ChIs from Sham-lesioned, 6-OHDA-lesioned, and 6-OHDA-lesioned mice treated with chronic LD. Scale bars are 1 sec and 50 pA. n=78-85 cells in each group. (C) Box and whiskers plots of spontaneous cell activity in the absence (79-85 cells in each group) and the presence (10-12 cells in each group) of synaptic blockers CNQX (10 µM), APV (25 µM) and picrotoxin (25 µM). (D) Coefficient of variation of instantaneous sAP frequencies in sham, 6-OHDA, and chronic LD groups (n=78-85). p<0.05 (*), p<0.01 (**), p<0.001 (***), or p<0.0001 (****) by Kruskal-Wallis test with Dunn’s multiple comparison test.

Next, using cell-attached recordings we measured spontaneous action potentials (sAP) of ChIs in acute striatal slices from the three groups of mice. Cholinergic neurons were identified by their large soma size relative to other cells in the dorsolateral striatum. The spontaneous firing frequency of ChIs decreased in 6-OHDA lesioned mice, but this decrease was reversed by chronic L-DOPA treatment to a level higher than that of sham-lesioned mice (**Figure 1B and 1C**). These changes persisted in the presence of blockers of ionotropic GABA and glutamate receptors (**Figure 1C**), indicating that altered spontaneous firing was driven by ChI-intrinsic mechanisms.

ChIs exhibit cell autonomous firing patterns, including irregular firing with bursts and pauses (Goldberg and Wilson, 2005; Wilson, 2005; Wilson and Goldberg, 2006). Previous reports showed that DA depletion decreased the regularity of spontaneous firing and the number of bursts and pauses (McKinley et al., 2019). Consistent with these findings, we found an increase in the coefficient of variation (CV) of the instantaneous frequencies of sAP in ChIs from 6-OHDA mice, which was restored to sham control levels after chronic L-DOPA (**Figure 1D**). Similarly, we found that 6-OHDA lesion decreased the number of both bursts and pauses, while chronic L-DOPA treatment of lesioned mice increased the number of bursts and restored the number of pauses compared to sham control levels. The duration of each burst was not different between the three groups, whereas pause duration was longer in the 6-OHDA and shorter in the chronic-LD groups. When the number of bursts and pauses were normalized to the number of sAP in each group, there was no significant difference between groups, suggesting that these irregularities of firing pattern changed in parallel with altered firing rate (**Figure S2A-F**). Thus, DA depletion significantly decreases spontaneous firing rate and regularity of ChI spiking, while chronic treatment with L-DOPA restores the key parameters of regularity while increasing firing frequency beyond that of control animals.

### Electrophysiological characteristics and synaptic input to striatal ChIs

To determine whether changes in basic electrophysiological characteristics of ChIs could explain observed alterations in spontaneous activity of the cells, we performed whole-cell recordings. Voltage-current dependence (**Figure 2A**), resting membrane potential (**Figure 2B**), input resistance (**Figure 2C**), membrane capacitance (**Figure 2D**), and rheobase (**Figure 2E**) were not different between the experimental groups. Similarly, analysis of the waveforms of spontaneous (**Figure 2F-H**) or evoked (**Fig 2I, J**) action potentials revealed no differences in peak amplitude, half width, rise time, decay time, threshold, and latency to first spike. However, current injections elicited fewer action potentials in ChIs from 6-OHDA lesioned mice indicating decreased excitability, which was reversed by chronic L-DOPA treatment (**Fig 2I, K**).

**Fig 2.**
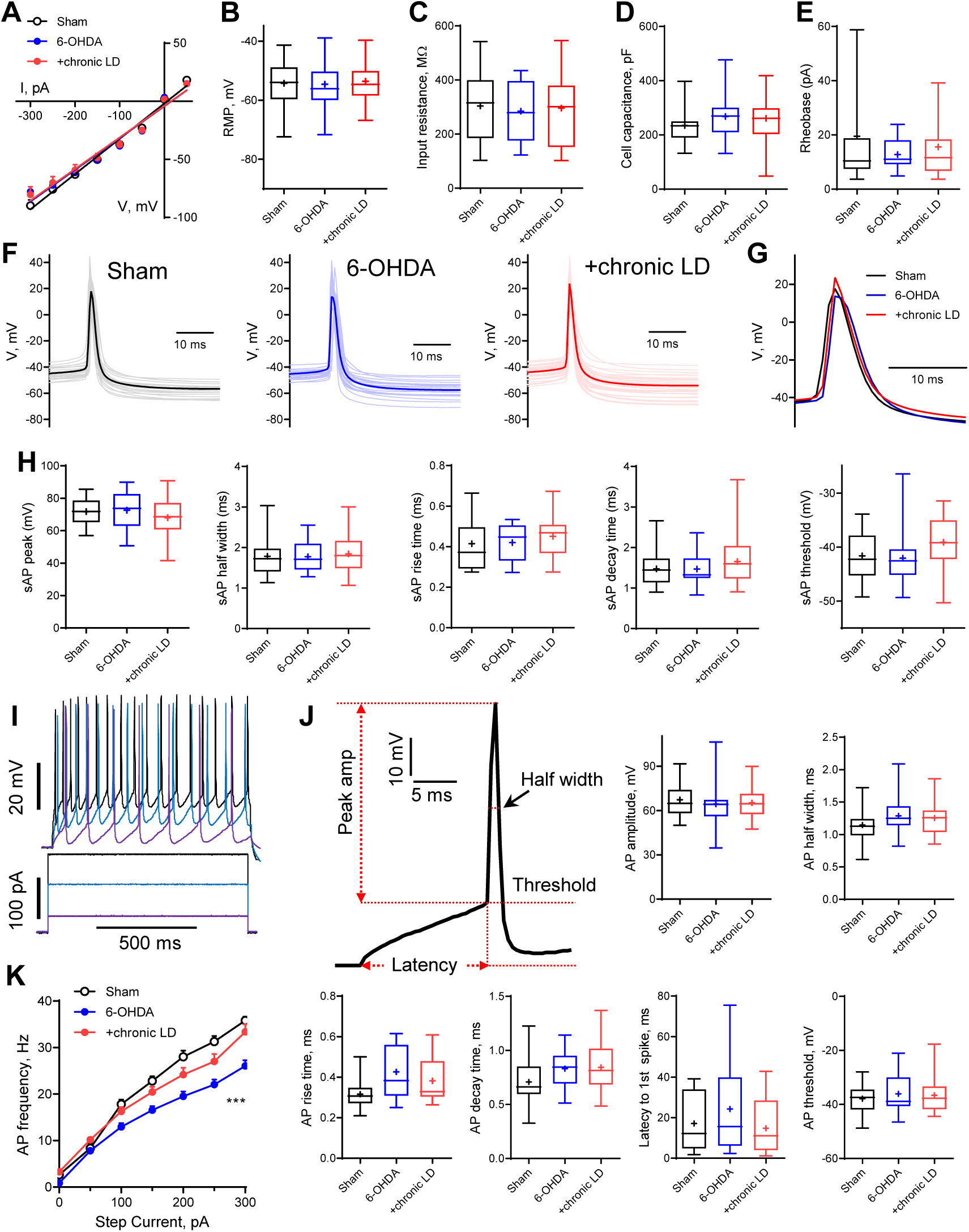
Basic electrophysiological characteristics in ChIs from control, 6-OHDA, and chronic LD mice. (A-E) Voltage-current dependence (A), resting membrane potential (B), input resistance (C), membrane capacitance (D) and rheobase (E) were unchanged in 6-OHDA and chronic L-DOPA (LD) groups. Recordings were conducted in the whole-cell mode in the presence of 1 µM TTX (n=7-34 neurons per group). (F-G) Representative (F) and averaged (G) traces of sAPs from the three experimental groups. (H) Shape characteristics of sAPs were not different between the groups (n=22-25 cells per group). (I) Representative traces of action potentials evoked by current injections. (J) Shape characteristics of action potentials evoked by 100 pA current injection were not different between the groups (n=18-28 cells in each group). (K) Number of evoked action potentials following current injection was decreased in ChIs from 6-OHDA lesioned mice but restored after chronic L-DOPA treatment. p<0.001 (***) from two other groups by mixed-effect analysis with Tukey’s post-hoc test (n=11-18 cells in each group).

Several reports have shown that changes in neuronal excitability correlate with remodeling of their dendritic arbors (Al-Muhtasib et al., 2018; Cazorla et al., 2012; Fieblinger et al., 2018). We assessed this with Sholl analysis of three dimensional neuritic arbor reconstructions from streptavidin labeled ChIs that were filled with biotin during recording (**Figure 3A**). ChIs from both the 6-OHDA and chronic-LD groups had more intersections with Sholl radii as a function of distance from the soma and more total intersections (**Figure 3B, D**). This was accompanied by greater total neurite length and a larger ending radius (**Figure 3C, E**), although the number of primary branches and ramification index were not different (**Figure 3F, G**). These data suggest that dopaminergic lesion induced a remodeling of ChI neurites that was not reversed with chronic L-DOPA treatment.

**Fig 3.**
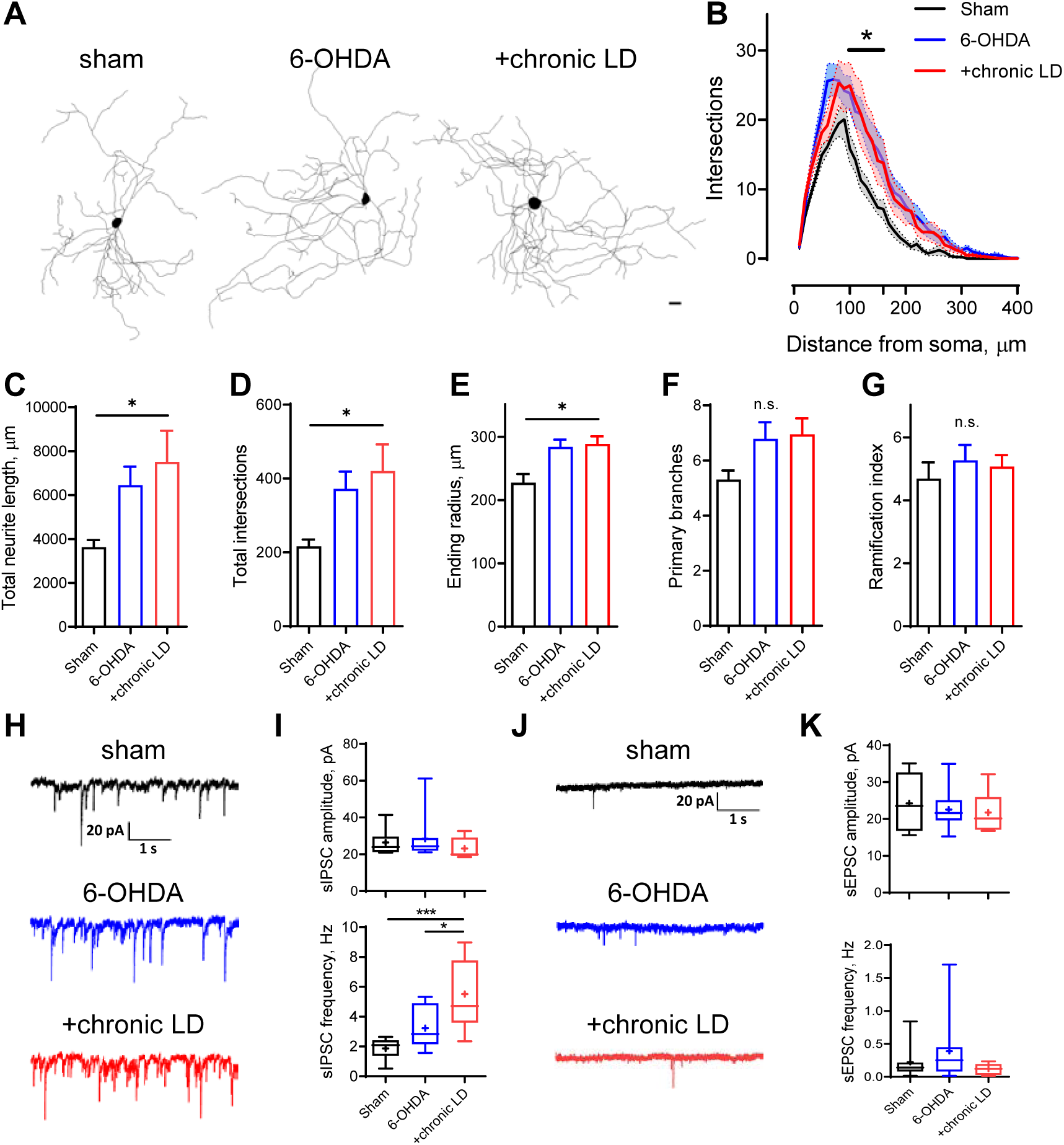
Morphological parameters and synaptic inputs in ChIs from control, 6-OHDA lesioned and chronic LD mice. (A-G) Sholl analysis of ChI morphology. Cells were filled with biocytin during physiological recordings, fixed, and imaged for biocytin labeling by confocal microscopy. (A) Representative maximum projection images from three dimensional reconstructions of ChIs, scale bar = 20 μm. (B) Sholl analysis of reconstructed ChIs, solid line denotes mean intersections at indicated distance from soma, shaded area = SEM, p<0.05 (*) for Sham vs. 6-OHDA and Sham vs. +chronic LD, Dunnett’s multiple comparison test following mixed-design ANOVA, p=0.0003 for interaction between treatment group and distance from soma. (C-G) DA lesion caused significant increase in total dendrite length (C), total number of intersections (D) and ending radius (E), whereas the number of primary dendrites (F) and ramification index (G) were similar between the groups, p<0.05 vs. Sham, Kruskal-Wallis test with Dunn’s multiple comparison (n=13-28 cells). (H and J) Representative traces of spontaneous inhibitory postsynaptic currents (sIPSCs) and excitatory postsynaptic currents (sEPSCs) of ChIs from sham, 6-OHDA and chronic LD mice. (I) Amplitudes of sIPSCs were similar, but their frequency was increased in chronic L-DOPA group. p<0.05 (*) and 0.001 (***) by one-way ANOVA and Tukey’s multiple comparisons test (n=7-10 cells). (K) No changes in amplitude or frequency of sEPSCs were detected (n=8-15 cells).

We next investigated whether changes in excitatory and inhibitory synaptic inputs followed observed adaptations in dendritic morphology. Spontaneous inhibitory postsynaptic currents (sIPSC) frequency was increased in the chronic-LD group compared to sham mice, while sIPSC amplitude was unchanged (**Figure 3H, I**). In contrast, neither frequency nor amplitude of spontaneous excitatory postsynaptic currents (sEPSC) was changed by the treatments (**Figure 3J, K**). These results demonstrate that L-DOPA treatment of DA lesioned mice increased synaptic connectivity of ChIs with GABA neurons, despite dendritic remodeling in both DA lesioned and chronic-LD groups.

### Restoration of DA depletion-induced decrease of HCN current by chronic L-DOPA

Examination of the whole-cell recording traces from ChIs of the three groups revealed marked differences in the kinetics of voltage change following the action potential. In the 6-OHDA group, the cell returned to the threshold firing potential at a significantly slower rate, while in the chronic-LD group, the hyperpolarization that followed the AP appeared to be lower in amplitude (**Figure 4A**). This prompted us to next look at the activity of the channels known to mediate the afterhyperpolarization (AHP) currents that regulate the rate of spontaneous firing of striatal ChIs.

**Fig 4.**
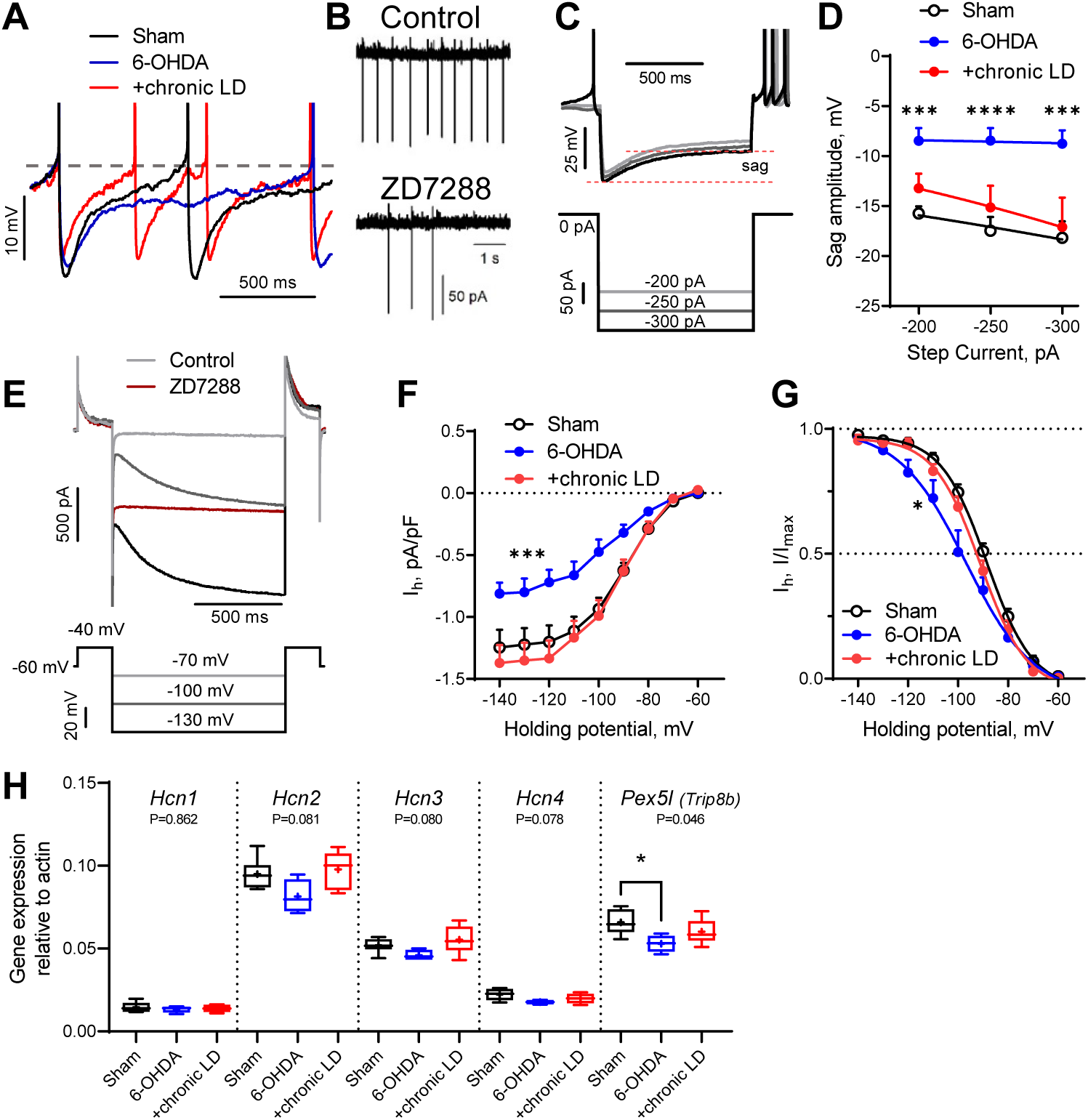
HCN-mediated currents are decreased by 6-OHDA lesion. (A) Representative perforated-patch recordings of sAP from Sham, 6-OHDA and chronic-LD groups. Traces were chosen to start at the same RMP. Dotted line represents threshold potential, which was similar between the groups. Note the markedly slower rate of cell depolarization after the action potential in the ChIs from 6-OHDA group. (B) Representative cell-attached recordings of ChI activity before and after treatment with HCN channel blocker ZD7288 (25 µM). (C) Current-clamp recordings showing voltage sag, a characteristic of HCN channel activation. (D) Quantification of sag amplitude at different current steps. p<0.01 (**) or 0.001 (***) compared to sham by two-way ANOVA with Tukey’s multiple comparison test (n=17-20 cells). (E) Voltage-clamp protocol (lower) and representative ZD7288-sensitive (*I*_*h*_) current (upper). (F) *I*_*h*_ density was decreased in ChIs from DA-depleted mice. (G) Boltzmann fits of normalized *I*_*h*_ densities. For F and G: p<0.05 (*) or p<0.001 (***) vs. both Sham and chronic LD groups by repeated measures two-way ANOVA with Tukey’s multiple comparisons test, n=10-16 cells. (H) ChI-specific gene expression of *Hcn1-4* isoforms and *Pex5l (Trip8b)* was measured by RT-qPCR from striatal mRNA immunoprecipitated from *ChAT-Cre:Ribotag* mice treated as indicated. Target mRNA levels were normalized to β-actin. P-values are for Kruskal-Wallis test, p<0.05 (*) with Dunn’s multiple comparisons test (n=4-6).

Hyperpolarization-activated cyclic nucleotide-gated (HCN) channels are essential for the cell-autonomous spontaneous firing of ChIs (Ferreira et al., 2014; Oswald et al., 2009; Wilson, 2005). Accordingly, bath application of an HCN channel antagonist ZD7288 (25 μM) significantly decreased ChI sAP frequency in our slice preparation (**Figure 4B**). To characterize HCN channel activity, we measured the characteristic voltage sag induced by hyperpolarizing current injection in the current clamp mode (**Figure 4C**) and also directly isolated ZD7288 sensitive HCN currents (*I*_*h*_) in the voltage clamp mode (**Figure 4E**). ChIs from 6-OHDA mice exhibited smaller voltage sag amplitudes and decreased *I*_*h*_ than either sham or chronic-LD mice, whereas there was no difference in these parameters in ChIs from chronic-LD mice compared to Sham (**Figure 4D, F)**. Furthermore, Boltzmann fits of normalized voltage dependences of HCN currents showed a more negative voltage threshold in ChIs from 6-OHDA mice compared to sham and chronic-LD animals (**Figure 4G**; V_50_ was −88.5±0.9 mV for Sham, −96.9±2.8 mV for 6-OHDA and −91.6±1.1 for chronic-LD group), suggesting that the gating properties or the expression profile of HCN isoforms were altered by 6-OHDA lesion and restored by chronic L-DOPA exposure (Simeone et al., 2005; Wang et al., 2001; Zolles et al., 2006).

We therefore next assessed whether the ChI-specific gene expression profiles of HCN channel isoforms were changed by DA depletion or chronic L-DOPA treatment using bitransgenic *ChAT-Cre:Ribotag* mice which express a tagged ribosomal subunit only in cholinergic neurons (Sanz et al., 2009). We subjected these mice to the same DA depletion and chronic L-DOPA treatment paradigm and then collected tagged ribosome-bound mRNA for RT-qPCR analysis. mRNA was harvested 20 hours after the last injection of saline or L-DOPA, reflecting steady-state gene expression levels. This technique yielded ∼38-fold enrichment of cholinergic neuron specific genes compared to total striatal mRNA (*Chat*:β-*actin*; ribotag: 0.423 ± 0.014 vs. input: 0.011± 0.001). Consistent with previous findings, *Hcn2* was the most abundantly expressed HCN isoform in ChIs (**Figure 4H**) while *Hcn3* and *Hcn4* mRNA were enriched in ChIs compared to total striatal input (not shown). There was no difference in mRNA levels of *Hcn1-4* between any groups, though there was a trend towards decreased levels of *Hcn2, Hcn3*, and *Hcn4* in the 6-OHDA group (**Figure 4H**). The trafficking and gating of HCN channels is regulated by tetratricopeptide repeat-containing Rab8b-interacting protein (TRIP8b; gene name *Pex5l*), an auxiliary β subunit expressed in neurons (Bankston et al., 2012; Lewis et al., 2011). TRIP8b mRNA levels were significantly reduced in ChIs from 6-OHDA mice (**Figure 4H**), which may contribute to the decrease in their HCN current. Together, these findings indicate that the decrease in firing rate of ChIs from 6-OHDA mice could be due to decreased HCN function. However, as HCN activity and expression of its subunits was restored to sham levels in the chronic-LD group, changes in *I*_*h*_ alone cannot account for the accompanying overshoot in ChI firing caused by chronic L-DOPA treatment.

### Persistent decrease in medium afterhyperpolarization currents

The AHP following each ChI action potential is mediated by potassium channels that are activated during cell depolarization and concomitant Ca^2+^ influx (Bennett et al., 2000; Goldberg and Wilson, 2005; Tubert et al., 2016). At low frequency spiking of ChIs, such as during their cell-autonomous activity, small conductance calcium-activated potassium channels (SK) mediate a medium-duration afterhyperpolarization (mAHP). Conversely, prolonged high frequency firing of ChIs activates a slow afterhyperpolarization (sAHP) that can last several seconds (Goldberg et al., 2009; Wilson and Goldberg, 2006).

To assess changes in AHP currents, first we applied a 300 ms depolarizing pulse from −60 mV to 10 mV, which induced a train of action potentials that was followed by a characteristic tail current lasting seconds after the end of the voltage pulse (**Figure 5A**). We measured the current during the *peak* response, which occurred within the first 200 ms following the offset of the voltage pulse, and during the long tail at 1000 ms (*late* response) (Bennett et al., 2000; Sanchez et al., 2011; Wilson and Goldberg, 2006). Both *peak* and *late* currents were inhibited by the non-selective K^+^ channel blocker barium (200 µM) (**Figure 5A**), though their amplitudes were unchanged in ChIs from 6-OHDA and chronic-LD mice compared to the sham group (**Figure 5B**). Next, to assess the currents that mediate mAHP more selectively, we used a shorter depolarization protocol, stepping from −60 mV to 0 mV for 100 ms (**Figure 5C**). As SK channels are the main mediators of mAHP currents, we used apamin (100 nM) to selectively block this channel (Goldberg and Wilson, 2005; Wilson and Goldberg, 2006). Apamin decreased *peak* AHP current without affecting the *late* component and revealed that SK channel activity was significantly decreased in ChIs from both 6-OHDA and chronic-LD groups compared to sham (**Figure 5D**).

**Fig 5.**
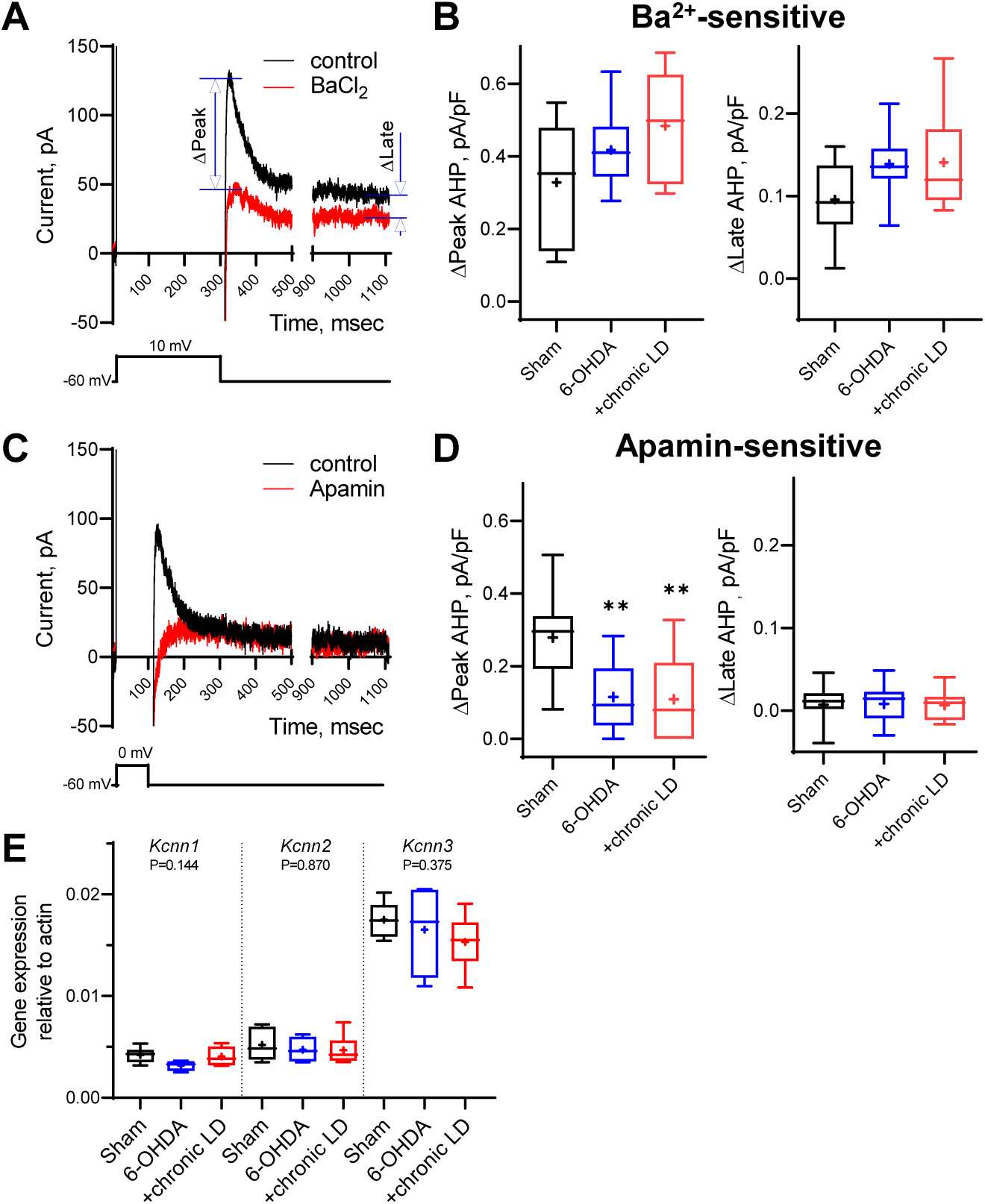
Changes in medium and slow afterhyperpolarization (AHP) currents. (A and C) Representative voltage-clamp recording of (A) Ba^2+^-sensitive and (C) apamin-sensitive AHP currents before and after drug application (upper traces), and corresponding voltage protocols (lower traces). (B) BaCl_2_ (200 µM) blocked both *peak* and *late* phases of the current but the magnitude of the changes was similar in all groups. (D) Using a depolarization protocol to recruit primarily mAHP currents, apamin (100 nM) decreased the *peak* current amplitude without altering the *late* stage of the AHP current. Apamin-sensitive currents were significantly decreased in 6-OHDA and chronic LD groups. p<0.01 (**), Kruskal-Wallis test with Dunn’s multiple comparison, n=13-15 neurons. (E) ChI-specific gene expression of *Kcnn1-3* (SK1-3) isoforms was measured by RT-qPCR as on Figure 4H. Target mRNA levels were normalized to β-actin. No significant differences between the groups by Kruskal-Wallis non-parametric ANOVA (n=4-6).

To determine whether changes in mAHP current were due to decreased expression of SK channels, we measured mRNA levels of SK1-3 (gene name *Kcnn1-3*) in ChIs from *ChAT-Cre:Ribotag* mice. There was no difference in the mRNA expression of these SK isoforms, suggesting that changes in mAHP current were not mediated by transcriptional regulation of these channels (**Figure 5E**).

### Partial inhibition of HCN and SK channels recapitulates changes in ChI firing rate caused by DA depletion and chronic L-DOPA treatment

Based on the changes in *I*_*h*_ and mAHP currents presented above, we hypothesized that altered activity of HCN and SK channels might be responsible for decreased spontaneous activity in 6-OHDA and increased spontaneous activity in chronic-LD mice (**Figure 6A**). To model these changes pharmacologically, we first established the concentrations of ZD7288 and apamin that provide ∼50% inhibition of HCN- and SK-mediated currents observed in ChIs from 6-OHDA and chronic-LD mice (**Figure 6B, C**). Next, using slice preparations from control mice (no sham surgery), we performed cell-attached recordings of ChI activity before and after bath application of 1 nM apamin either alone (to mimic decreased SK but normal HCN activity seen in chronic-LD group) or in combination with 1 µM ZD7288 (to mimic inhibition of both SK and HCN channels in 6-OHDA group). Partial blockade of both channels decreased sAP frequency (**Figure 6D, E**), increased the coefficient of variation of sAP frequency (**Figure 6F, G**) and phenocopied changes in bursting and pausing of ChIs in the 6-OHDA group (**Figure S2G-L**). Similarly, the features of ChI firing in the chronic-LD group were recapitulated with partial SK current blockade, including increased sAP frequency (**Figure 6D, E**), increased coefficient of variation of sAP frequency (**Figure 6F, G**) and similar changes in the bursting/pausing patterns (**Figure S2**). Overall, our findings suggest that alterations in ChI spontaneous activity caused by DA depletion and subsequent chronic L-DOPA treatment can be reproduced by decreasing HCN- and SK-mediated currents.

**Fig 6.**
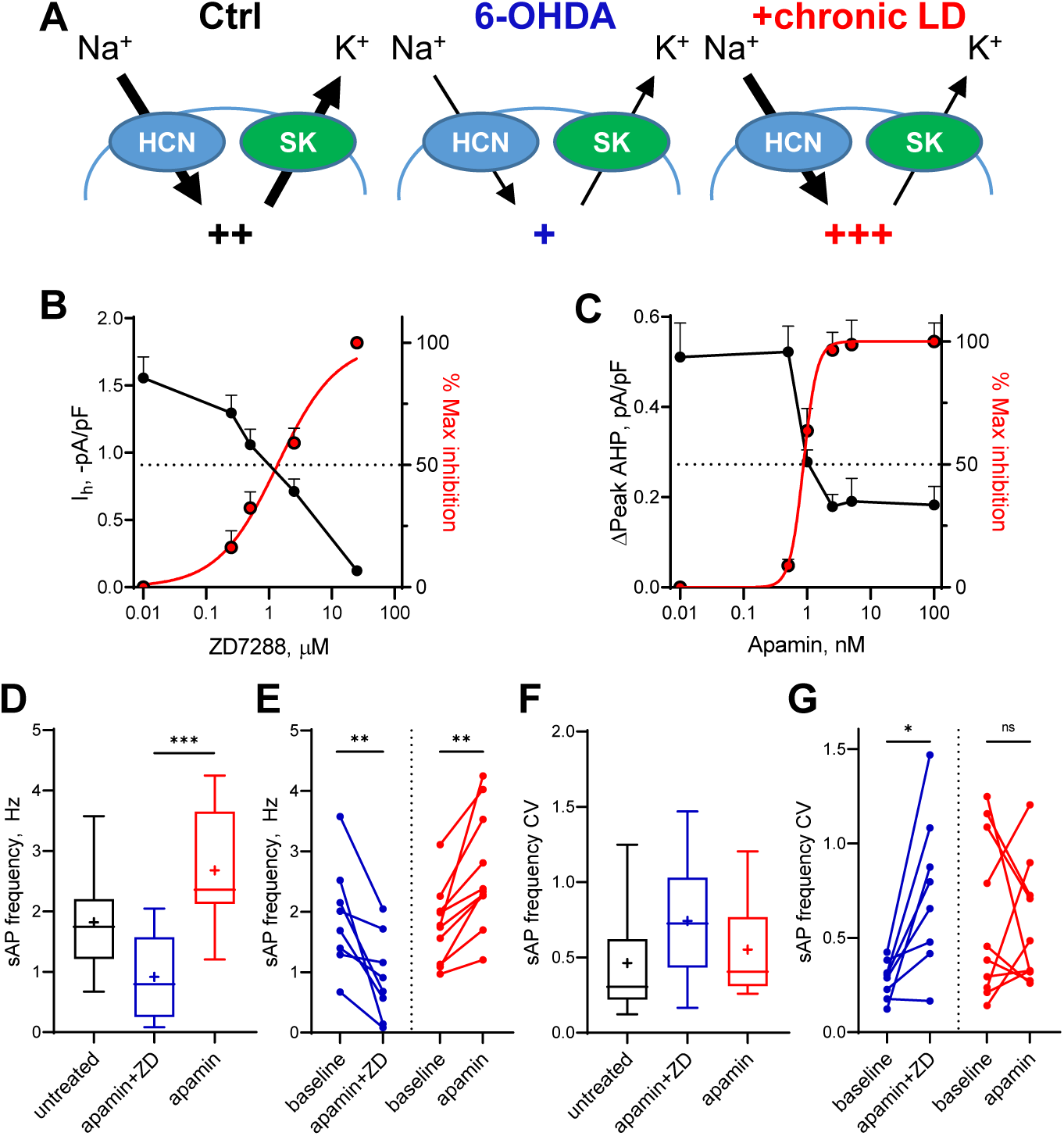
Partial inhibition of HCN and SK channels is sufficient to mimic changes in ChI activity after lesion and chronic L-DOPA treatment. (A) Proposed changes in HCN and SK currents in ChIs from different treatment groups and their effect on spontaneous firing rates (+). (B) Dependence of *I*_*h*_ density (as measured on Fig 4E) on ZD7288 concentration. Red curve represents fit of the data with the equation Y=100(X^Slope^)/(IC_50_^Slope^ + X^Slope^). Hill slope and IC_50_ are 1 and 1.4 µM, correspondingly. (C) Dependence of mAHP current density (as measured on Figure 5C) on apamin concentration. Hill slope and IC_50_ are 4 and 0.9 nM, correspondingly. (D-G) Partial SK and HCN channel blockade reproduced changes in the sAP rate and coefficient of variation of 6-OHDA and chronic LD groups. Although on average the decrease and increase in sAP frequency caused by apamin+ZD and apamin alone, respectively, did not reach statistical significance (D), apamin+ZD reliably decreased and apamin alone increased baseline sAP firing rate in individual ChIs (E). Likewise, the differences in the median coefficient of variation did not reach statistical significance (F), however, apamin+ZD increased CV over baseline for most ChIs, whereas apamin alone did not change the baseline CV (G). p=0.0006 (*** in panel D) by Kruskal-Wallis non-parametric test with Dunn’s multiple comparisons post-hoc analysis. In panels E and G, p<0.05 (*) and p<0.01 (**), Wilcoxon matched-pairs signed rank test (n=8-17).

### Changes in DA receptor sensitivity

DA receptor-mediated signaling has been shown to be altered in ChIs in the DA depleted striatum (Araki et al., 2000; Ding et al., 2006; Zhou et al., 2014). We thus measured the response of ChIs to bath application of 30 µM DA. Similar to Pitx3 mutant mice (Ding et al., 2011), both naïve and L-DOPA treated 6-OHDA lesioned mice showed enhanced sensitivity of ChIs to DA exposure (**Figure S3**). In the presence of D1/D5 receptor antagonist SCH 23390 (thus revealing the contribution of inhibitory D2/D3 receptors) DA slightly decreased spontaneous firing in all groups, although the effect did not reach statistical significance. Likewise, application of D2/D3 agonist quinpirole trended to decrease the firing rates in all groups also without reaching statistical significance. Pre-treatment with a combination of SCH 23390 and D2/D3 antagonist sulpiride abolished the effect of DA on sAPs, confirming that the effects were DA receptor-mediated. We therefore conclude that DA signaling undergoes significant alteration from balanced inhibitory/excitatory in intact animals to mostly D1/D5 excitatory in both 6-OHDA and chronic-LD groups (**Figure S3D**). Together with altered firing activity of ChIs, these changes in responsiveness to DA receptor ligand binding could contribute to the manifestation of LID.

## Discussion

Dysregulation of cholinergic neurotransmission is an important contributor to both the expression of PD symptoms and the adverse effects of DA replacement therapy. Here, we examined the consequences of DA loss and those of chronic treatment with L-DOPA on mouse ChI physiology. In addition to altered morphology, synaptic connectivity and responsiveness to DA receptor stimulation, we found significant alterations of cell-intrinsic properties of ChIs. In the DA depleted striatum, both HCN- and SK-mediated currents were diminished. HCN current reduction was accompanied by reduced TRIP8b, which regulates trafficking and gating of HCN channels, rather than changes in HCN mRNA expression. Interestingly, chronic treatment of lesioned mice with L-DOPA restored HCN activity to sham levels while SK currents remained depressed. The pharmacological blockade of HCN and SK channels to mimic the DA depleted and chronic L-DOPA treated states recapitulated the changes in rate and pattern of ChI firing, providing new insights into molecular adaptations that follow striatal DA depletion in PD patients receiving L-DOPA therapy.

### Changes in ChI tonic activity following DA depletion and chronic L-DOPA treatment

Anticholinergic drugs have been effective in the treatment of PD symptoms, highlighting the importance of cholinergic neurotransmission to basal ganglia function. Early observations showed that the severity of PD symptoms in patients is worsened by drugs that increase cholinergic activity (Duvoisin, 1967), while more recent reports demonstrated that direct optogenetic and chemogenetic inhibition of ChIs improved DA depletion-mediated motor dysfunction (Maurice et al., 2015; Tanimura et al., 2019; Ztaou et al., 2016). Though the contribution of aberrant ChI neurotransmission to PD pathophysiology is unquestioned, conflicting results on the change in ChI activity caused by DA depletion have been reported. We show here that both spontaneous activity and excitability of dorsolateral striatal ChIs in 6-OHDA injected mice are significantly decreased. Similar changes were recently demonstrated using a genetic model with diphtheria toxin to induce dysfunction in dopaminergic neurons (McKinley et al., 2019). In contrast, other studies using similar protocols in rodents have shown no significant change in the tonic firing rate (Ding et al., 2006), and increased excitability of ChIs (Maurice et al., 2015; Sanchez et al., 2011; Tubert et al., 2016). Some of these discrepancies can be attributed to differences in animal species (mice vs. rats) or experimental conditions (DA-lesion protocol, composition of the recording solutions, etc.) but careful evaluation of the contribution of these differences is currently lacking.

Levels of acetylcholine in striatal tissue or CSF from PD patient samples are not different from controls (Duvoisin and Dettbarn, 1967; Welch et al., 1976), but these approaches cannot resolve differences in acetylcholine release due to altered synaptic activity. The lack of change in overall acetylcholine levels raises the question of whether the “hypercholinergic state” in PD reflects an increase in basal cholinergic release, a shift in the balance between DA and acetylcholine levels (McKinley et al., 2019), or altered firing of ChIs in response to physiological stimuli. For example, striatal tonically-active neurons (putative ChIs) in non-human primates exhibit a pause response that is acquired after training in a classical conditioning task (Aosaki et al., 1994). In trained animals, the conditioned stimulus elicits a brief pause (∼200 ms) in ChI tonic firing, followed by a brief rebound increase in firing rate. Lesion of the striatonigral system with 1-methyl-4-phenyl-1,2,3,6-tetrahydropyridine (MPTP) did not change the tonic firing rate of ChIs, but abolished this pause response *in vivo* (Aosaki et al., 1994). As this pause is thought to provide a “window” to encode cortico-striatal plasticity (Deffains and Bergman, 2015), loss of this pause response and increased variability of ChI firing (Figure 1D) (McKinley et al., 2019) may cause dysregulation of cortico-striatal neurotransmission and aberrant striatal plasticity. Additionally, the loss of DA may change the response of striatal neurons to acetylcholine as the degree of muscarinic and nicotinic acetylcholine receptor binding is altered in the brain tissue of PD patients (Aubert et al., 1992; Joyce, 1993; Pimlott et al., 2004) and after experimental DA depletion (Cremer et al., 2015; Joyce, 1991). Combined with the dendritic remodeling observed in our results (Figure 3) and other studies (Lozovaya et al., 2018), and changes in the connectivity from ChIs to spiny projection neurons (Salin et al., 2009) that may drive changes in the regulation of neurotransmitter release from striatonigral synapses (Borgkvist et al., 2015), it is likely that the regulation of striatal output by ChIs is perturbed as a result.

Chronic L-DOPA treatment of 6-OHDA-lesioned mice increased the tonic firing of ChIs beyond the sham lesioned group (Figure 1), while restoring many firing pattern abnormalities (Figure S2) and neuronal excitability (Figure 2K) back to sham levels. This may represent the cellular mechanism underlying the hypercholinergic state that contributes to motor dysfunction in PD patients who have undergone DA replacement therapy. Consistent with this idea, ablation of ChIs reduces LID (Won et al., 2014) and direct activation of ChIs increases LID (Aldrin-Kirk et al., 2018; Bordia et al., 2016a) in rodent PD models, although the effect of ChI modulation is complex (Bordia et al., 2016a; Divito et al., 2015; Maurice et al., 2015). In aggregate, these studies suggest that there is an interaction between an increased rate and improper pattern/context of ChI firing in PD and after L-DOPA treatment. Reinstating both of these aspects of cholinergic neurotransmission may be critical for restoring the functional behavior.

In slice preparations, ChIs fire spontaneously and generate a variety of spiking patterns and firing frequencies autonomously. Though excitatory and inhibitory inputs to ChIs are present, these are largely silent as the firing rate and pattern of ChIs are unaffected by blockade of AMPA, NMDA, GABA_A_, D1, D2, or muscarinic receptors (Bennett and Wilson, 1999). Consistent with this, the changes in tonic firing rate caused by DA depletion and chronic L-DOPA treatment persisted in the presence of blockers of ionotropic glutamate and GABA receptors (Figure 1C), indicating that these changes are intrinsic to ChIs. Furthermore, the increases in dendritic branching and GABAergic inputs onto ChIs after chronic L-DOPA treatment (Figure 3), which likely represent compensatory remodeling of striatal circuitry, did not have significant effect on the tonic firing of ChIs in our preparation. Together with altered responsiveness to DA receptor stimulation (Figure S3), these synaptic adaptations may, however, play important roles *in vivo*, which will be addressed in future studies.

### DA depletion and chronic L-DOPA treatment change HCN activity

HCN channels are essential for the tonic firing of ChIs and serve to depolarize neurons back towards spike threshold during the hyperpolarization that follows an action potential. Inhibition of *I*_*h*_ decreases the rate and increases irregularity of tonic firing in ChIs (Bennett et al., 2000; Deng et al., 2007; McKinley et al., 2019; Zhao et al., 2016). In line with this, we found that DA depletion reduced *I*_*h*_ and shifted the gating for channel activation to more negative potentials (Figure 4), which was accompanied by slower and irregular firing. By contrast, chronic treatment of DA depleted mice with L-DOPA restored *I*_*h*_ to the level of sham-lesioned animals, though the firing rate exceeded sham levels. These findings are consistent with a recent study using a targeted diphtheria toxin model of DA depletion which found a similar decrease of *I*_*h*_ that was mediated by the transcriptional downregulation of specific *Hcn* isofroms (McKinley et al., 2019). Although we also observed a trend for decreased HCN2-4 mRNA, the differences did not reach statistical significance (Figure 4H). In contrast, we found that DA depletion significantly decreased Trip8b mRNA levels in ChIs. Trip8b is expressed in neurons and regulates surface expression and trafficking of HCN channels. Knockdown of Trip8b *in vivo* causes the mis-localization of HCN channels and reduces *I*_*h*_ in hippocampal CA1 neurons. Our results confirm a previous report of reduction in Trip8b mRNA after DA depletion in the external globus pallidus (GPe) in conjunction with decreased *I*_*h*_ (Chan et al., 2011). Chronic treatment with L-DOPA reversed these trends in gene expression of Trip8b and HCN channels, suggesting that striatal DA, including the levels achieved with once daily administration of L-DOPA, maintains HCN activity. This effect lasts longer than pharmacokinetic availability of L-DOPA (Abercrombie et al., 1990) since the recordings were performed at least 20 hours after the last L-DOPA dose, reflecting the long-term adaptation to chronic L-DOPA exposure rather than the immediate pharmacologic actions of L-DOPA itself. A possible mechanism linking DA depletion with changes in HCN biophysical properties may involve the regulation of cAMP production by DA receptor signaling (Greengard, 2001). Binding of cAMP to HCN channels, which can also be regulated by Trip8b (Hu et al., 2013; Saponaro et al., 2014), shifts their activation kinetics towards more positive potentials and increases channel opening kinetics (Wainger et al., 2001). We found that HCN activation was shifted to more hyperpolarized potentials in ChIs from DA depleted mice and restored by chronic L-DOPA treatment (Figure 4G), although whether these changes are mediated by decreased cAMP availability or other DA-dependent mechanisms remains to be tested.

### DA depletion decreases SK current

During tonic firing of ChIs, calcium entry following each action potential induces outward potassium current mediated by apamin-sensitive SK-channels (Goldberg and Wilson, 2005) and the kinetics of this afterhyperpolarization current determines the firing rate of ChIs (Bennett et al., 2000). We found decreased SK current in ChIs from DA depleted mice both with or without L-DOPA treatment, consistent with smaller AHP amplitudes. Although the reduction in SK current was the same for both groups, only parkinsonian mice treated chronically with L-DOPA showed higher firing rates, indicating that in the 6-OHDA group the decrease in firing rate from HCN current loss cannot be overcome by the increase in firing rate expected to result from reduction in SK current. The mRNA levels of SK channel isoforms were not different between groups indicating that the changes in channel activity are mediated by posttranscriptional mechanisms. As SK channel activation is coupled to calcium entry through Ca_v_2.2 (N-type) channels in ChIs, the reduction in SK channel activity could indicate persistent dysregulation of these calcium channels (Goldberg and Wilson, 2005). Interestingly, M4 muscarinic autoreceptor-activated calcium currents through Cav2.1/2.2 channels are decreased following DA depletion (Ding et al., 2006), although no persistent decrease in SK activity has been reported. Whether decreased SK channel activity is caused by deficient auto-receptor signaling remains to be tested, although no effect of muscarinic antagonists on spontaneous firing of ChIs was previously found (Bennett and Wilson, 1999).

### Pharmacological blockade of HCN and SK channels in ChIs mimics firing patterns caused by DA depletion and chronic L-DOPA treatment

Several changes in spontaneous firing of ChIs in our animal models were recapitulated by pharmacological inhibition of HCN and SK channels in striatal slices from control animals, including altered spontaneous activity and rhythmicity. Critically, we were able to titrate the concentrations of antagonists to resemble the degree of channel inhibition caused by DA depletion and chronic L-DOPA treatment. Consistent with previous studies, a saturating concentration of apamin caused ChIs to transition into burst firing mode, characterized by increased firing frequency within bursts, long pauses between bursts and decreased average firing frequency (data not shown and (Bennett et al., 2000; Yorgason et al., 2017)). However, when SK channels were only partially inhibited using an IC_50_ concentration of apamin, the average firing rate of ChIs was increased without an increase in burstiness, closely mimicking changes observed in the ChIs from L-DOPA treated mice. Likewise, partial blockade of both HCN and SK channels caused significant depression of ChI firing rate, consistent with the DA depleted state. These findings confirm that reducing the current from these channels is sufficient to account for the changes in ChI activity and show that the slower rate of depolarization from hyperpolarized potentials caused by HCN inhibition overrides the smaller AHP resulting from reduced SK activity. Importantly, further studies should address whether targeted restoration of these channels’ activities can be employed to ameliorate PD-related motor deficiencies, such as akinesia and LID.

In summary, we followed up on our previous finding that chronic L-DOPA treatment of DA lesioned mice is associated with increased ChI firing and now demonstrate a novel cellular mechanism that implicates changes in HCN and SK currents that results from DA depletion and chronic L-DOPA treatment. Critically, these changes are evident beyond the pharmacological time course of L-DOPA, suggesting that they reflect the physiological state of ChIs upon which DA replacement works in patients treated with L-DOPA. Interestingly, we found that only some of the changes in ChI physiology caused by DA depletion are restored by chronic L-DOPA treatment and some persisted despite treatment. Our data suggest that HCN and SK channels can be targeted for therapeutic intervention, although development of cell type-specific modulators of channel activities is desired for effective and safe behavioral outcomes.

## Acknowledgements

This work was supported by NINDS R01NS101982 (UJK), NINDS R01NS075222 (EVM), the JBP Foundation (DS) and NIDA R0107418 (DS). We thank Nicolas Tritsch for helpful comments and Joanna Garcia for technical assistance.

## Author Contribution

Conceptualization, YD, EVM, UJK; methodology, YD, SJC, TCM; investigation, SJC, TCM, YD, TC, NJ, EVM; writing, TCM, EVM, UJK; revisions, SJC, TCM, YD, TC, DS, EVM, UJK; visualization, TCM, EVM; supervision, EVM, DS, UJK; and funding acquisition, UJK

## Declaration of Interest

none

## Star Methods

### Animals

The use of the animals followed the National Institutes of Health guidelines and was approved by the Institutional Animal Care and Use Committee of Columbia University and New York State Psychiatric Institute. For behavior and slice electrophysiology studies, we used male C57BL/6J mice (Jackson Laboratory, Bar Harbor ME, stock# 000664) at twelve-weeks of age at the beginning of the experiments. To obtain bitransgenic mice that express “tagged” ribosomes selectively in cholinergic neurons, mice expressing the Cre-recombinase under the regulation of the ChAT promotor (ChAT-Cre; Tg(ChAT-Cre)GM24sat/Mmucd from the GENSAT Project obtained from the MMRRC, stock# 017269-UCD) (Gong et al., 2007) were bred with mice expressing a Cre-activated knock-in of HA-tagged *RPL22* ribosomal subunit (Ribotag mice; B6N.129-Rpl22(tm1.1Psam)/J obtained from Jackson Laboratory, stock# 011029) (Sanz et al., 2009). Mice of both sexes were used for experiments and were homozygous for Ribotag and heterozygous for ChAT-Cre.

### DA lesion and chronic L-DOPA treatments

Anesthesia was induced by intraperitoneal (IP) injection of ketamine and xylazine, followed by a subcutaneous injection of bupivacaine for local anesthesia at the incision site. Animal were head-fixed in a stereotaxic apparatus (Kopf Instruments) with ear cups, and 6-hydroxydopamine (6-OHDA, Sigma H-116; 4.5 ug dissolved in 1.5 microliters of 0.05% ascorbic acid in 0.9% saline) was injected into left medial forebrain bundle (MFB) (coordinates: AP −1.3 mm and ML +1.3 mm from Bregma, and DV −5.4 mm from skull surface) through a small borehole in the skull. The entire volume was infused over 7.5 minutes through a stainless-steel cannula (Braintree Scientific, RM-SBL STD), which was left in place for an additional 5 min before withdrawal and incision closure. Desipramine (Sigma D3900; 25 mg/kg delivered IP) was given 30 min prior to 6-OHDA infusion to block uptake of the toxin by noradrenergic neurons. Intensive post-operative care included providing supplemental nutrition (Bacon Softies F-3580, Bio-Serv) and extra fluids (saline subcutaneously and dextrose saline IP). The health status of the animals was monitored daily until stabilization of body weight with free access to food and water. Sham-lesioned control mice received same volume of vehicle into the left MFB.

3-4 weeks after unilateral 6-OHDA injection, lesioned mice were tested for stepping (Fig. S1A) and randomly divided to receive daily IP injections of saline or L-DOPA (3 mg/kg + 12.5 mg/kg benserazide), while all sham-injected control mice received saline. For slice electrophysiology studies, mice were used at 3-11 weeks after the first injection of saline or L-DOPA. For gene expression analysis, striatal tissue was collected after 3 weeks of daily L-DOPA treatment. Dopaminergic lesion was confirmed in some cohorts of mice by western blot analysis of striatal tissue lysates for tyrosine hydroxylase (TH) protein levels which showed near complete depletion of TH in the striatal hemisphere ipsilateral to the 6-OHDA lesion (0.0184 ± 0.003-fold contralateral hemisphere; not shown).

### Weight-supported treadmill stepping task

Stepping tests to assess akinesia of the impaired forelimb (contralateral to the 6-OHDA lesion) were performed 3-4 weeks post 6-OHDA injection before the initiation of repeated L-DOPA or saline treatment. Both forelimbs were placed on a treadmill with the surface moving at 4.6 cm/sec away from the head of the mouse while the body weight was supported by examiner. Forepaw steps were video recorded from five nonconsecutive cycles of treadmill. The number of left and right paw steps was counted over a distance of 45 cm per trial for five non-consecutive trials and averaged to obtain the stepping score per mouse.

### LID assessment

L-DOPA-induced abnormal involuntary movements (AIMs) were assessed after the first IP injection of L-DOPA (the acute L-DOPA in naïve state) and again after 3 weeks of daily L-DOPA treatment (chronically-treated state) (Fig. S1B-D). One cohort of mice was tested again at 10 weeks of L-DOPA treatment, which showed no difference from the 3 week time point indicating that LID scores remained stable throughout the time course of the experiments (not shown). At the start of the session, each mouse was placed into a clear polypropylene cylinder and allowed to acclimate for at least 3 min. L-DOPA was then injected and the mouse was video-recorded for 1 min periods at 0, 5, 10, 20, 40, 60, 80, 100 and 120 min post injection. Limb and axial dyskinesias were analyzed from recorded videos using previously described protocols (Ding et al., 2011; Won et al., 2014).

### Slice preparation and electrophysiological recordings

At the time of recordings, mice were 4-6-month-old; each day, animals were randomly selected from a different treatment group. Slices were prepared at least 20h after the last L-DOPA or vehicle treatment. Mice were euthanized by cervical dislocation and coronal 270 µm-thick striatal slices were prepared on a vibratome (VT1200; Leica, Sloms, Germany) in oxygenated ice cold cutting-artificial cerebrospinal fluid (ACSF) containing (in mM): 194 sucrose, 30 NaCl, 4.5 KCl, 26 NaHCO_3_, 6 MgCl_2_·6H_2_O, 1.2 NaH_2_PO_4_, and 10 D-glucose (pH 7.4, 290 ± 5 mOsm). Slices were then transferred to oxygenated normal ACSF containing 125.2 NaCl, 2.5 KCl, 26 NaHCO_3_, 1.3 MgCl_2_·6H_2_O, 2.4 CaCl_2_, 0.3 NaH_2_PO_4_, 0.3 KH_2_PO_4_, and 10 D-glucose (pH 7.4, 290 ± 5 mOsm) at 34 °C and allowed to recover for at least 40 min before the recordings.

Electrophysiological recordings were performed on an upright Olympus BX50WI (Olympus, Tokyo, Japan) microscope equipped with a 40x water immersion objective, differential interference contrast (DIC) optics and an infrared video camera. All recorded ChIs were located in the dorsolateral striatum and identified by larger somatic size than neighboring neurons. Slices were transferred to a recording chamber and maintained under perfusion with normal ACSF (1.5-2 mL/min) at 34°C. Patch pipettes (3-5 MΩ) were pulled using P-97 puller (Sutter instruments, Novato, CA) and filled with internal solutions as indicated below. All chemicals for ACSF as well as gramicidin, biocytin, dopamine, picrotoxin, sulpiride, and SCH23390 were purchased from Sigma (St. Louis, MO). TTX, apamin, ZD7288, APV, and CNQX were from Tocris (Bristol, UK). Patch clamp recordings were performed with a MultiClamp 700B amplifier (Molecular Devices, Forster City, CA) and digitized at 10 kHz with InstruTECH ITC-18 (HEKA, Holliston, MA). Data was acquired using WINWCP software (developed by John Dempster, University of Strathclyde, UK) and analyzed using Clampfit (Molecular Devices), Igor Pro (Wavemetrics, Lake Oswego, OR), and Matlab (MathWorks, Natick, MA).

The pipette solution contained (in mM): 115 K-gluconate, 10 HEPES, 2 MgCl_2_, 20 KCl, 2 MgATP, 1 Na_2_-ATP, and 0.3 GTP (pH = 7.3; 280 ± 5 mOsm). After measuring cell firing rate in cell-attached mode, the neuronal membrane was ruptured and basal electrophysiological characteristics of ChIs, including IV curve, RMP, input resistance and membrane capacitance were measured in current-clamp mode. To measure rheobase, a 75 pA/sec current ramp from - 50 to +100 pA was applied (Maurice et al., 2004). For perforated-patch recordings, 60 mg/ml gramicidin was added to the patch pipette solution.

Glutamatergic spontaneous excitatory postsynaptic currents (sEPSCs) were recorded in whole-cell voltage clamp mode in cells pre-treated with 25 µM picrotoxin to inhibit GABA_A_ receptors; the internal pipette solution contained (in mM) 120 CsMeSO_3_, 5 NaCl, 10 HEPES, 1.1 EGTA, 2 Mg^2+^-ATP, 0.3 Na-GTP, 2 Na-ATP, and 5 QX314 to block of voltage-activated Na^+^ channels (pH = 7.3, 280 ± 5 mOsm). GABAergic spontaneous inhibitory postsynaptic currents (sIPSCs) were recorded in the presence of 25 µM APV, and 10 µM CNQX (or NBQX) to inhibit glutamatergic receptors; internal pipette solution contained (in mM): 140 CsCl_2_, 2 MgCl_2_, 10 HEPES, 2 EGTA, 2 MgATP, 1 Na_2_-ATP, 0.3 GTP and 5 QX314 (pH = 7.3, 280 ± 5 mOsm). Postsynaptic currents were detected and analyzed using Mini Analysis program (Synaptosoft, Decatur, GA). The threshold for amplitude detection was 8-10 pA which was >2-fold the RMS of the background noise.

Voltage sag was measured in current-clamp mode following 500ms-long current injections from 0 to −300 pA. To measure *I*_*h*_ currents in voltage-clamp mode in the presence of TTX, ChIs were held at −60 mV, then depolarized to −40 mV followed by 1s hyperpolarizing steps to −140 mV in 10 mV voltage increments. Sensitivity to 25 µM ZD7288 was used to confirm that the voltage sag and *I*_*h*_ were mediated by HCN channels.

To measure Ba^2+^-sensitive afterhyperpolarization (AHP) currents, cells were clamped at −60 mV in the presence of TTX followed by a 300ms-long depolarizing step to +10 mV. *Peak* current amplitude was measured in the first 200ms following the offset of the depolarizing voltage step, while the *late* phase was the average steady-state current at 900-1000 ms after the end of the step. To measure apamin-sensitive current, cells were depolarized from −60mV to 0 mV for 100ms. BaCl_2_ (200 µM) or apamin (0.5 – 100 nM) were applied for 10 min and currents measured after drug application were subtracted from those before the treatments to assess the contribution of *peak* and *late* components of AHP.

### Analysis of Burst-Pause Activity

We used the Robust Gaussian Surprise (RGS) method to determine burst and pause patterns during tonic ChI firing using the MatLab code available from (Storey et al., 2016). This method identifies differences in the firing rate of adjacent spikes against the local log ISI distributions, assigning individual spikes to burst or pause strings based on statistical criteria in comparison to the Gaussian distribution of the entire spike train (Ko et al., 2012). The parameters used for burst and pause detection were: p=0.15 for the calculation of the central location, alpha=0.05 for Bonferroni correction, N_min_=2 for minimum number of spikes to be considered a burst/pause, and central distribution calculated as the median ± 2 x median average deviation.

### Immunohistochemistry and morphological analysis

For morphological characterization of recorded ChIs, biocytin (1 mg/ml) was included in the internal pipette solution and allowed to fill the cell for 30-40 min after achieving the whole-cell configuration. Then, slices were fixed with 4% paraformaldehyde in 0.1M PBS overnight at 4°C, washed with Tris-buffered saline (TBS: 50 mM Tris-Cl, 150 mM NaCl, pH 7.5) and incubated with a streptavidin-Dylight 633 conjugate (1:200; Thermo Scientific) in TBS + 0.6% Triton-X100 for 48h. The slices were then rinsed and mounted on glass slides using Fluormount-G (Southern Biotech). For morphological reconstruction, serial optical sections encompassing the neurites of biocytin-labeled ChIs were imaged at 0.25 μm^2^ pixels at a z-depth of 0.74 μm using a Leica DM6 confocal microscope with a 20x/0.7 NA oil immersion objective (Leica HCX PL APO CS). The neurites were traced from the resulting stacks using the Simple Neurite Tracer plugin in Fiji (ImageJ) for Sholl analysis (Ferreira et al., 2014; Longair et al., 2011).

### Gene expression analysis

Brain tissue from ChAT-Cre x Ribotag mice was collected at least 20 hours after the last L-DOPA or saline injection. Striatal tissue ipsilateral to the 6-OHDA lesion from 3-4 mice for each experimental group was pooled per replicate (4-6 total replicates per condition), weighed and immediately placed into cold homogenization buffer at 5% (w/v) consisting of (pH 7.4, in mM): 50 Tris, 100 KCl, 12 MgCl_2_, 1 DTT, 1% Nonidet P40 substitute (Roche), 0.1 mg/ml cyclohexamide, 1x protease inhibitor cocktail, and 200 U/mL RNAsin (Promega). The tissue was then homogenized with a powered Dounce homogenizer at 1700 rpm for 14 complete up-down strokes, followed by centrifugation at 10,000 RCF for 10 minutes at 4°C. 5 μl of mouse anti-HA antibody (HA.11, BioLegend MMS-101R) was added to 800 μl of the resulting supernatant and incubated for 4 hours at 4°C. The mixture was then added to protein-G magnetic beads (Dynabeads 400 μl equivalent, Invitrogen) and incubated overnight at 4°C. The next day, the beads were separated from the supernatant with a magnet and washed 3 times for 10 minutes at 4°C in high salt washing buffer consisting of (in mM): 50 Tris, 300 KCl, 12 MgCl_2_, 0.5 DTT, 1% Nonidet P40 substitute, and 0.1 mg/ml cyclohexamide. After the last wash, the beads were collected and the bound RNA eluted with 350 μl of RLT buffer from the RNeasy Micro Kit. The beads were removed from the RLT buffer and the RNA was isolated according the manufacturer’s instructions (Qiagen). The resulting RNA was assayed for integrity and amount with a Bioanalyzer (Agilent). All samples exhibited a RNA integrity number (RIN) score of 8.4-10.

cDNA libraries were generated from the resulting RNA with the SuperScript IV VILO master mix (Invitrogen) according to the manufacturer’s protocols. Gene expression was measured using Taqman chemistry with probes for the target genes and *Actb* as a housekeeping control assayed in duplex for each well on a CFX96 Touch Real-Time PCR Detection System (BioRad) using TaqMan Fast Advanced (Applied Biosystems) master mix according to the manufacturer’s instructions. Each sample was assayed in triplicate using 0.33 μl of the cDNA library (undiluted) per reaction. Cycling conditions were: 50° x 2 mins, 95° x 20s, (95° x 3s, 60° x 30s) x 40 cycles. Gene expression is expressed as a ratio of the target gene to *Actb* as determined by 2^-ΔCt^. The probes were obtained from Applied Biosystems and include: *Hcn1* (Mm00468832_m1), *Hcn2* (Mm00468538_m1), *Hcn3* (Mm01212852_m1), *Hcn4* (Mm01176086_m1), *Pex5l* (Mm00458088_m1), *Chat* (Mm01221882_m1), *Kcnn1* (Mm01349167_m1), *Kcnn2* (Mm00446514_m1), *Kcnn3* (Mm00446516_m1) and *Actb* (4352341E).

### Statistical analysis

Unless stated otherwise, electrophysiological data represent observations from single neurons in slices from an indicated number of animals. For data expressed as box and whisker plots, the whiskers denote the range of all data points, the box the 25-75th percentile, the horizontal line the median, and the cross the mean. For bar graphs, data are expressed as mean ± standard error of the mean. The statistical tests used are indicated in the figure legends and in the text. Statistical analysis and data were plotted using GraphPad Prism 8.0 (GraphPad Software, San Diego, CA).

## Supplemental information

**Fig S1.**
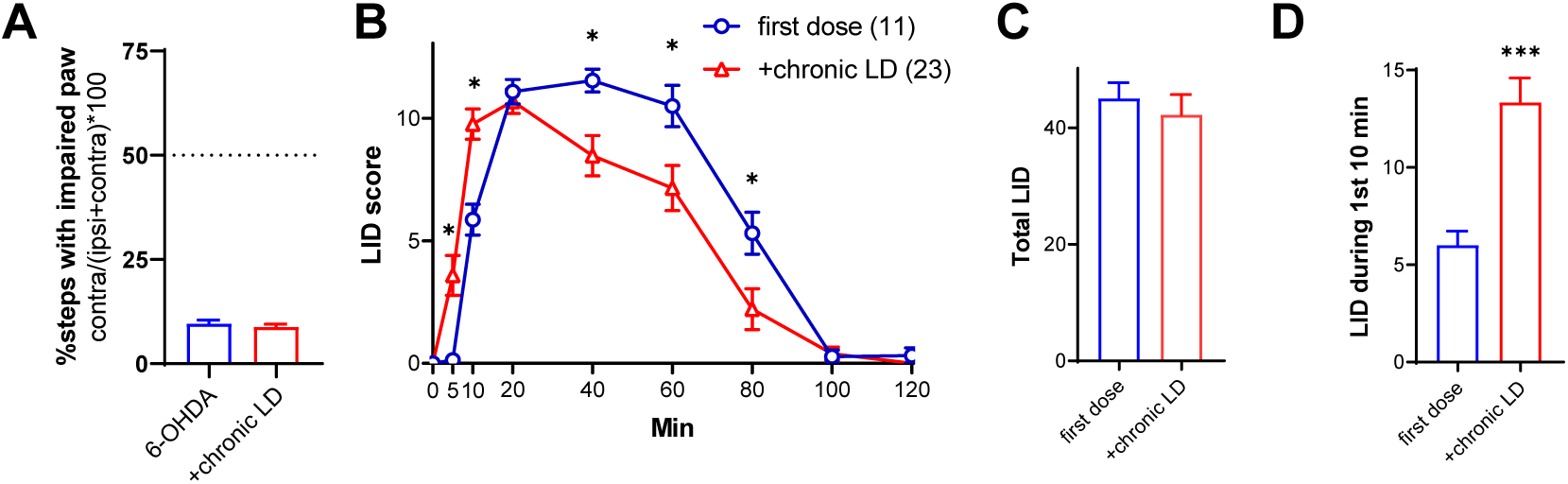
Expression of L-DOPA-induced dyskinesia (LID) in 6-OHDA-lesioned mice. (A) All 6-OHDA-lesioned mice developed contralateral front paw stepping deficits, p<0.0001 vs. 50% for both groups, Wilcoxon signed rank test. There was no difference in stepping deficit between the mice assigned to the 6-OHDA-only or chronic L-DOPA groups, p=0.5467 by Mann-Whitney test. (B) Time course of LID expression following single injection of L-DOPA (3 mg/kg) either as the first dose or after chronic administration of L-DOPA. p<0.0001 for time x treatment interaction by two-way ANOVA), (*) p<0.05 between first dose vs. chronic administration at the same time point after L-DOPA injection by Bonferroni post-hoc test. (C) Total LID was not different between the groups, p=0.2946 by Mann-Whitney test. (D) LID expression occurred sooner after L-DOPA injection in mice that received chronic L-DOPA, p=0.0002, Mann-Whitney test.

**Fig S2.**
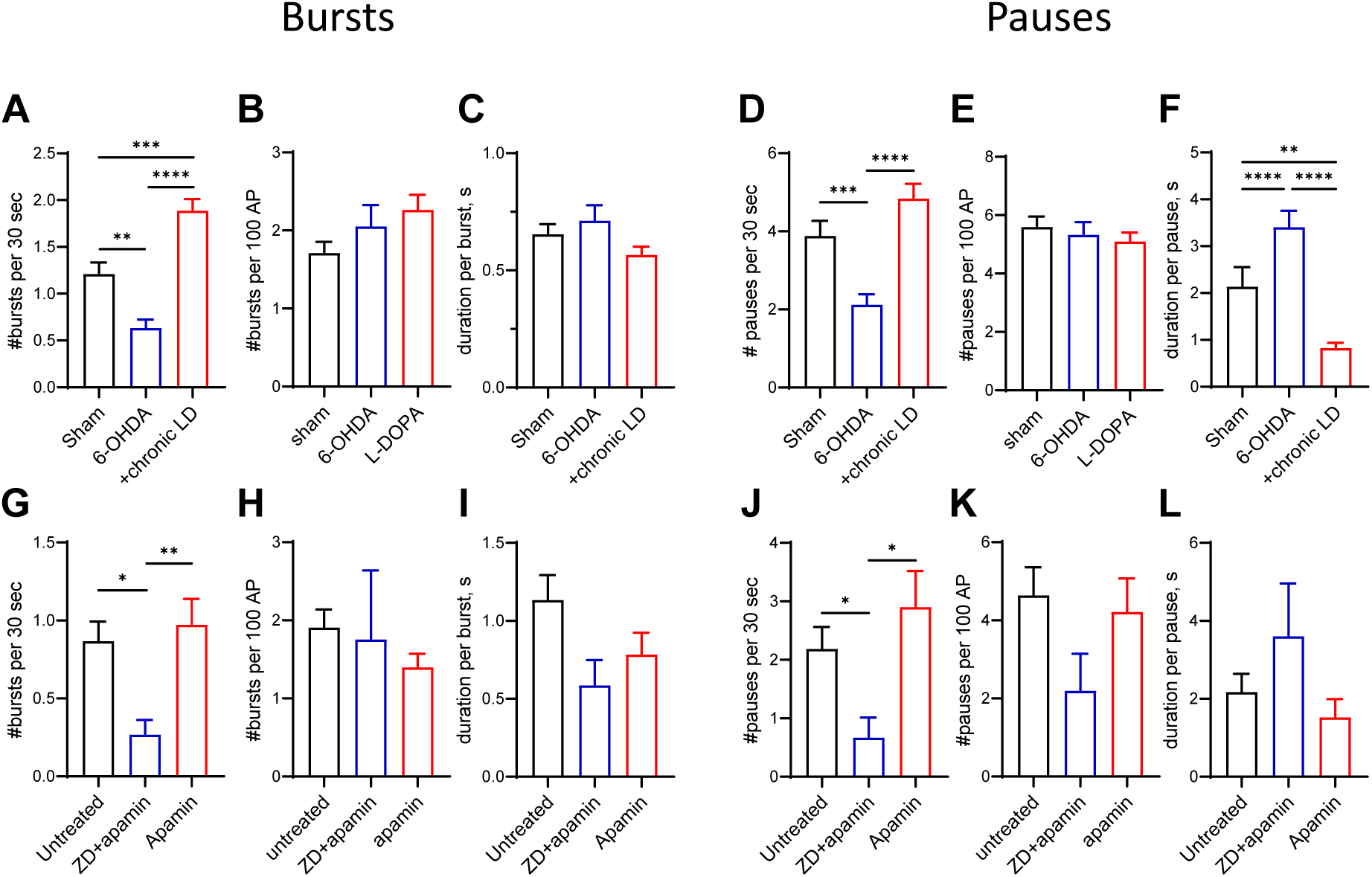
Burst-pause activity in ChIs. The number and duration of bursts and pauses were quantitated using the robust gaussian surprise method. (A-F) Firing patterns of ChIs from Sham, 6-OHDA, chronic LD groups. (G-L) Firing patterns of ChIs in control mouse brain slices treated with ZD7288 (1 µM) + apamin (1 nM) or the same concentration of apamin alone. Effect of treatments on (A,G) number of bursts per unit of time, (B,H) number of bursts normalized by the number of sAP, (C,I) burst duration, (D,J) number of pauses per unit of time, (E,K) number of pauses normalized by the number of sAP, and (F,L) pause duration. Blockade of HCN and SK currents largely reproduced changes in firing patterns observed in the 6-OHDA and chronic LD groups. Also note that number of bursts and pauses was not different between any groups when bursts or pauses were normalized to the number of action potentials (B,H,E,K), suggesting that the expression of bursts and pauses is related to the change in the firing rate. p<0.05 (*), p<0.01 (**), p<0.001 (***), p<0.0001 (****) Dunn’s multiple comparison test following Kruskal-Wallis nonparametric one-way ANOVA, n=83-87 neurons for A, B, D, and E; n=40-80 neurons for C and F; n=8-17 neurons for G, H, J and K; and n=5-17 for panel I and L.

**Fig S3.**
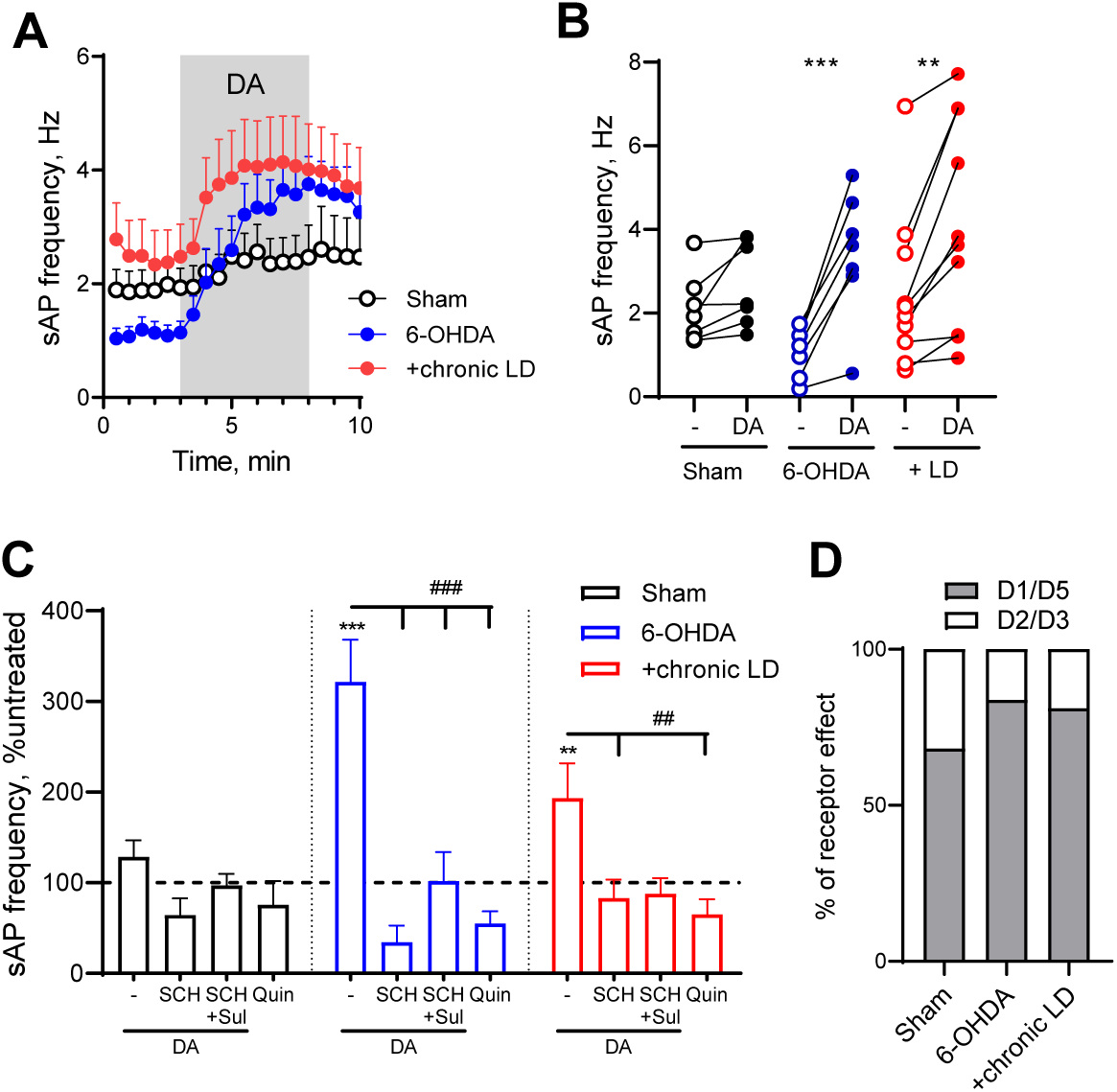
Altered DA receptor-mediated responses in ChIs from 6-OHDA lesioned mice before and after chronic L-DOPA treatment. (A) Perforated-patch recordings of sAP in ChIs following 30 µM DA perfusion in the presence of synaptic blockers. (B) Average sAP frequencies in individual cells before and after DA exposure. p<0.01 (**) or p<0.001 (***) by paired t-test. (N=8-12 cells in each group). (C) Normalized sAP frequencies in cells exposed to DA in the presence of synaptic blockers with or without D1/D5 receptor antagonist SCH23390 (10 µM) either alone or in combination with D2/D3 antagonist sulpiride (10 µM). Combination of SCH23390 and sulpiride completely abolished the effect of DA on sAPs in all groups. D2/D3 receptor agonist quinpirole (1 µM) had similar effect as a combination of DA and SCH23390. Dashed line represents sAP frequency in untreated cells. p<0.001 (***) vs. corresponding untreated controls or p<0.05 (#) or p<0.001 (###) by one-way ANOVA (N=5-8 cells in each group). (D) Relative contribution of excitatory (D1/D5) and inhibitory (D2/D3) receptor-mediated changes in sAP firing frequencies in response to DA treatment. Contribution of the D2/D3 signaling was calculated as a difference between untreated sAP firing frequency and that after ChI treatment with either quinpirole or a combination of DA and SCH23390. Contribution of D1/D5 receptors was estimated as a difference between sAP frequency in the presence of DA alone and DA with SCH23390. Groups are significantly different (p<0.0155) by chi-square test.

